# Decoys reveal the genetic and biochemical roles of redundant plant E3 ubiquitin ligases

**DOI:** 10.1101/115071

**Authors:** Chin-Mei Lee, Ann Feke, Christopher Adamchek, Kristofor Webb, José Pruneda-Paz, Eric J. Bennett, Steve A. Kay, Joshua M. Gendron

## Abstract

The ubiquitin proteasome system (UPS) is the main cellular route for protein degradation in plants and is important for a wide range of biological processes including daily and seasonal timing. The UPS relies on the action of E3 ubiquitin ligases to specifically recognize substrate proteins and facilitate their ubiquitylation. In plants, there are three major challenges that inhibit studies of E3 ligase function: 1) rampant genetic redundancy, 2) labile interactions between an E3 ligase and its cognate substrates, and 3) a lack of tools for rapid validation of bona fide substrates. To overcome these 3 challenges, we have developed a decoy method that allows for rapid genetic analysis of E3 ligases, *in vivo* identification of substrates using immunoprecipitation followed by mass spectrometry, and reconstitution of the ubiquitylation reaction in mammalian cells to rapidly validate potential substrates. We employ the strategy to study the plant F-box proteins, ZTL, LKP2, and FKF1 revealing differential genetic impacts on circadian clock period and seasonal flowering. We identify a group of circadian clock transcriptional regulators that interact with ZTL, LKP2, and FKF1 *in vivo* providing a host of potential substrates that have not been seen previously. We then validate one substrate of ZTL, the plant circadian clock transcription factor CHE, and show that ZTL mediates CHE ubiquitylation and that the levels of the CHE protein cycle in daily timecourses. This work further untangles the complicated genetic roles of this family of E3 ligases and suggests that ZTL is a master regulator of a diverse set of critical clock transcription factors. Furthermore, the method that is validated here can be tool employed widely to overcome traditional challenges in studying redundant plant E3 ubiquitin ligases.

## Introduction

The ubiquitin proteasome system (UPS) is an enzymatic pathway that mediates the covalent attachment of ubiquitin to substrate proteins(1, 2). The critical substrate targeting step of this cascade is mediated by E3 ubiquitin ligases(3, 4). E3 ligases must contain at least two protein-protein interaction domains: 1) a substrate recognition domain that interacts with specific substrate proteins and 2) a domain that recruits a ubiquitin-charged E2 enzyme either directly or as part of a larger protein complex. Together these domains act to bridge an interaction between the substrate and activated ubiquitin, catalyzing transfer of ubiquitin to lysine residues within substrate proteins.

The SKP1/CULLIN/F-BOX (SCF) complex is a multi-subunit E3 ubiquitin ligase complex that is conserved across eukaryotes(3, 4). Within this complex, F-box domain-containing proteins act as substrate adaptors using a highly diverse group of protein-protein interaction domains(5, 6). The F-box domain is an approximately 45 amino acid domain that binds SKP1 (7-9). SKP1 subsequently interacts with the cullin subunit that binds to a RING domain-containing protein (RBX1). RBX1 acts to recruit ubiquitin-charged and activated E2 enzymes to facilitate ubiquitylation of target proteins.

Identifying the functions of F-box proteins can be challenging for three specific reasons. First, in many species, there is widespread functional redundancy in the F-box family which makes traditional forward genetic studies difficult or impossible. This problem is greatly exacerbated in plants, as the *Arabidopsis thaliana* F-box gene family is one of the largest in the genome, containing nearly 700 members(6). Second, the interaction between an F-box protein and its substrate is often difficult to detect because the substrate protein is degraded or dissociates following ubiquitylation. Third, validating putative E3 ubiquitin ligase substrates can be challenging for the same reasons listed above.

These challenges are exemplified by a subfamily of three F-box proteins that regulates the circadian clock and seasonal flowering time in plants. The LOV/F-box/Kelch repeat family of F-boxes has three members: ZEITLUPE (ZTL), LOV KELCH PROTEIN 2 (LKP2), and FLAVIN-BINDING, KELCH REPEAT, F-BOX 1 (FKF1)(10-12). This F-box subfamily shares the same protein domain structure and has significant overlap in sequence identity. They contain a unique arrangement of protein interaction domains that allow them to communicate light signals to the circadian clock and factors controlling seasonal flowering time(13-17). The N-terminal LIGHT OXYGEN VOLTAGE (LOV) domain is a blue light photoreceptor that interacts with the regulatory protein GIGANTEA (GI) in a light-dependent manner(18-21). It is also predicted that the LOV domain of ZTL, LKP2, and FKF1 interact with the substrate proteins TIMING OF CAB2 EXPRESSION 1 (TOC1, sometimes called PRR1) and PSEUDO-RESPONSE REGULATOR 5 (PRR5), two homologous transcription factors that regulate the circadian clock(21-29). Furthermore, the FKF1 LOV domain interacts with CONSTANS (CO), a critical promoter of flowering(30, 31). Interestingly, TOC1 and PRR5 are destabilized through these interactions while CO is stabilized. The central region of the protein contains the F-box domain, which recruits the ubiquitylation machinery through binding to multiple variants of ARABIDOPSIS SKP1 HOMOLOGUE 1 (ASK1)(32-35).The C-terminal domain is a typical protein-protein interaction domain composed of Kelch repeats. It is predicted that both FKF1 and LKP2 interact with the CYCLING DOF FACTOR (CDF) family of flowering regulators through their Kelch repeat domains(14, 36, 37).

Genetic studies of the LOV/F-box/Kelch repeat gene family exemplify difficulties in studying redundant E3 ubiquitin ligases. Although ZTL, FKF1, and LKP2 play roles in the circadian clock and regulation of seasonal flowering time, their functions have diverged partially but not entirely, leading to complicated redundancy. For instance, the *ztl* knockout lengthens clock period by 4 hours while individual *lkp2* and *fkf1* knockouts have little or no effect on the clock(10-12). The *fkf1* knockout delays flowering time dramatically while *ztl* and *lkp2* knockouts have minimal effect on flowering(11, 12, 38, 39). Higher order loss-of-function mutants show additive effects, revealing roles for ZTL and LKP2 in photoperiodic flowering and LKP2 and FKF1 in circadian control(39, 40). Further complexity has been demonstrated by yeast two-hybrid experiments and co-immunoprecipitation in heterologous expression systems. These assays show that the clock factors TOC1 and PRR5 interact with all three family members(40), while the flowering regulator CDF proteins show binding preference for FKF1 and LKP2(14). These assays relied on prior knowledge of the substrates’ role in clock function to identify their relationship with the E3 ligases. To our knowledge no unbiased approaches have been used to discover additional putative substrates of this family of F-box proteins.

A variety of proteomics techniques can be used to identify the substrates of E3 ubiquitin ligases(41). These include substrate trapping, proximity labeling, enrichment using ubiquitin remnants, affinity purification techniques, and shotgun proteomics [Reviewed in(42)]. Furthermore, genetic techniques have been used to overcome functional redundancy including knocking out entire gene families(40), overexpressing full length E3 ubiquitin ligases(12), or expressing dominant-negative E3 ubiquitin ligases that prevent interaction with the ubiquitylation machinery(43-45).

We have developed a streamlined three-step method that couples dominant-negative genetic analysis, *in vivo* substrate identification, and rapid heterologous substrate validation for studying the function of plant F-box proteins or other E3 ubiquitin ligases. This includes engineering affinity-tagged dominant-negative F-box proteins and expressing them in Arabidopsis. This allows for analysis of the genetic contribution of an F-box gene and substrate identification using immunoprecipitation followed by mass spectrometry in the first transformant generation. We have named this the “decoy” method because the F-box deletion proteins “lure” the targets from the endogenous F-box proteins, trapping the interaction and allowing for genetic and biochemical analyses. We then validate putative substrates by reconstituting the ubiquitylation reaction for plant F-box proteins in the mammalian tissue culture system.

We tested the genetic effects of expressing ZTL, LKP2, and FKF1 decoys on the circadian clock and flowering in Arabidopsis. We show that ZTL has the greatest effect on the circadian clock, followed by LKP2 then FKF1. This relationship is inverted with respect to flowering time, as the FKF1 decoy has a strong effect on flowering while LKP2 and ZTL have weaker effects. These genetic results are corroborated by IP-MS studies, showing that FKF1 decoys interact with flowering regulators while LKP2 and ZTL decoys interact with clock factors. We further employ this unbiased target identification method to identify CCA1 HIKING EXPEDITION (CHE), a critical clock regulator(46), as a putative target of ZTL. We validate CHE as a ZTL target by reconstituting ZTL ubiquitylation of CHE in mammalian tissue culture cells and show that in plants the CHE protein cycles and is degraded in the dark, as would be expected of a ZTL substrate. The results validate a robust three step approach that can identify the genetic and biochemical function of F-box-containing E3 ubiquitin ligases in plants and other systems.

## Results

### Genetic and developmental analyses of ZTL, FKF1, and LKP2 decoy transgenic lines

Functional redundancy and the labile nature of F-box/target interactions complicate functional characterization of F-box proteins, particularly in plants. To bypass traditional challenges, we developed a “decoy” method to reveal the genetic contributions of F-box genes and provide a streamlined method of target identification and validation. In brief, the substrate recognition domain of an F-box protein is expressed without the F-box domain (Fig. 1a), allowing it to interact with targets but preventing recruitment of the machinery to ubiquitylate the substrate(43, 44, 47-49). Genetically, the decoy functions in a dominant-negative manner by competitively inhibiting ubiquitylation of substrates by any endogenous F-box proteins. Previously point mutants in the F-box domain of ZTL were suggested to retain some interaction with ASK proteins in Arabidopsis(33), making it important to delete the F-box domain completely to achieve a dominant-negative genetic effect. In addition, a dual affinity tag (3XFLAG 6XHis) allows for immunoprecipitation followed by mass spectrometry (IP-MS) approaches to detect stabilized candidate target proteins (Fig. 1a)(50). Thus, the decoy method couples previously developed genetic and biochemical techniques, allowing us to test the genetic roles of any plant F-box gene or gene family and then rapidly identify interacting proteins, thereby overcoming two of the major challenges in studying F-box proteins.

**Figure 1.**
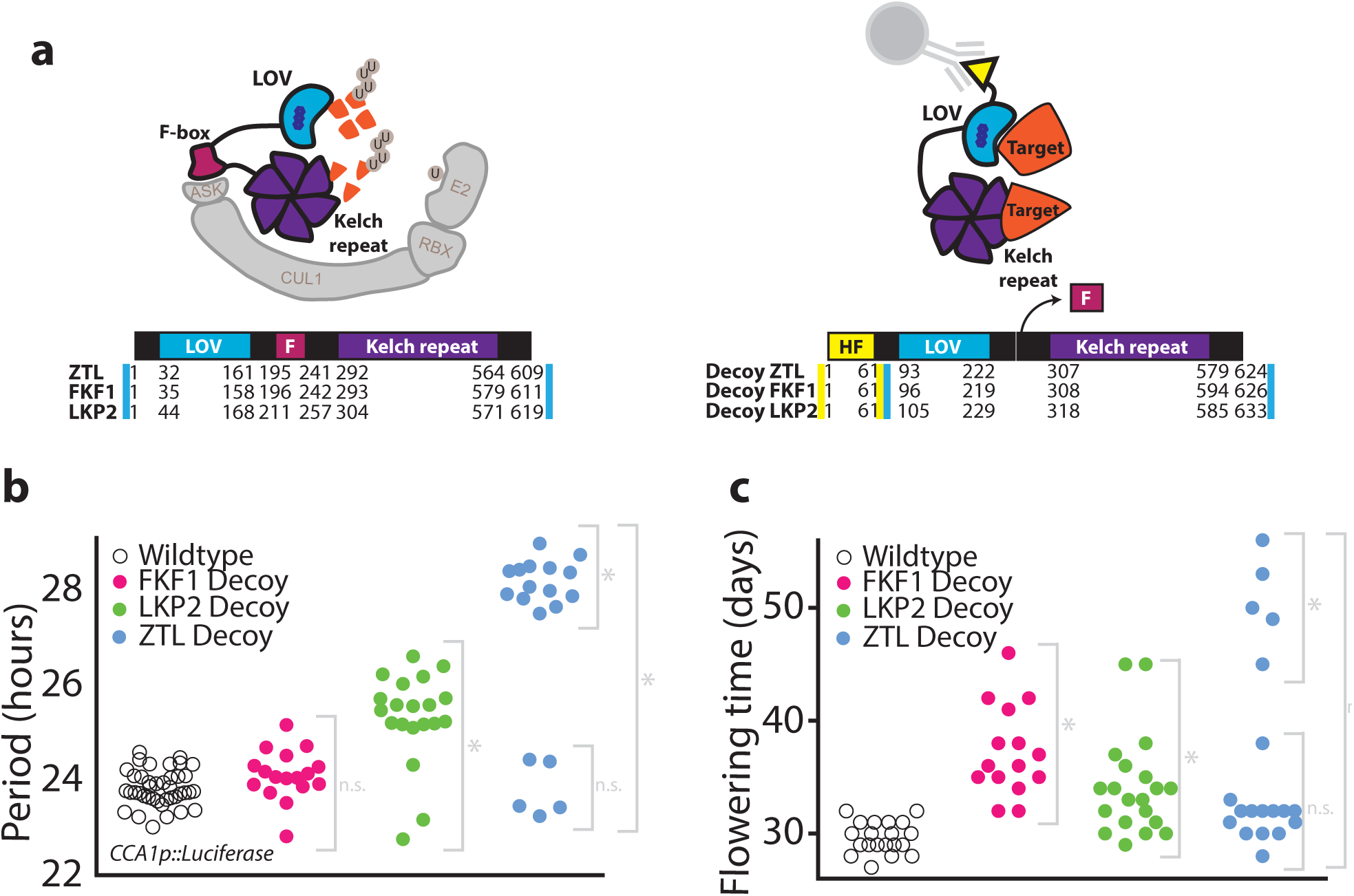
Decoys reveal the genetic contributions of redundant F-box genes. (a) Schematic of the decoy strategy applied to LOV-Kelch domain containing F-box proteins from Arabidopsis. F-box proteins bind to substrates and ubiquitylate them. This occurs through interaction with E2 ubiquitin ligases to facilitate proteasomal destruction. Decoy proteins lacking the F-box domain interact with their substrates but cannot target them for degradation, and the stabilized decoy-substrate complex can be immunoprecipitated through the His-FLAG tag. Below, amino acid numbers are shown for the three domains of ZTL, LKP2, and FKF1 wildtype and decoy proteins. (b) Period lengths (as measured by *CCA1p::Luciferase* activity) for T1 transgenic decoys from ZTL (blue), LKP2 (green), FKF1 (pink), and the wildtype parental line (*CCA1p::Luciferase* - open circles). (c) Flowering time (number of days for inflorescence stem to reach 1cm) for T1 transgenic decoys from ZTL (blue), LKP2 (green), FKF1 (pink), and the wildtype parental line *(CCA1p::Luciferase* - open circles). Grey brackets define individual groups used for statistical testing against the wildtype control using a Welch’s t-test with a Bonferroni-corrected α of 1.67×10^−3^. ^*^ represents p < α, n.s p > α. HF = His-FLAG. F = F-box.

ZTL, LKP2, and FKF1 compose a small family of partially redundant F-box proteins that control the circadian clock and seasonal flowering time. We sought to clarify their genetic relationship using the decoy method. To do this, we removed the F-box domain from each, leaving the LOV and Kelch repeat domains fused to one another, and added an affinity tag (Fig. 1a). We transformed *ZTL, LKP2,* and *FKF1* decoys into Arabidopsis harboring a circadian clock reporter transgene *[CIRCADIAN CLOCK ASSOCIATED 1 (CCA1)* promoter driving the expression of *Luciferase, CCA1p::Luciferase*]. We entrained the plants in 12 hour light/12 hour dark (LD) conditions for 7 days, and then transferred them to constant light (LL) for one day prior to imaging in LL. We performed two biological replicates of imaging and flowering time experiments, each with tens of T1 transformants, to avoid artifacts arising from expression variation from genomic insertion site.

ZTL decoy lines have a circadian period of approximately 28 hours and differ significantly from the wildtype period average of slightly less than 24 hours (p < .001) (Fig. 1b, S1, S2, and table S1). Furthermore, all ZTL decoy transgenics maintained rhythmicity, unlike overexpression of ZTL with point mutations in the F-box domain(33). Expression of the FKF1 decoy has the least effect on the circadian clock of the three (Fig. 1b, S1, S2, and table S1). The longer period in the FKF1 decoy transgenic line is statistically different than wildtype when data from multiple experiments is summed (p < .001), but is not always statistically significant when experiments are considered individually (Table S1). The LKP2 decoy lines cluster around a 26 hour period and differ significantly from wildtype(p < .001) (Fig. 1b, S1, S2, and table S1). FKF1 and ZTL decoys match period defects seen in knockout mutants(10, 11). Interestingly, LKP2 is expressed ubiquitously at lower levels than ZTL, the *lkp2* knockout has no effect on period, and *LKP2* overexpression causes arrhythmicity(12). Expression of the LKP2 decoy has a 2 hour lengthening of period which has not been observed in the previous genetic analyses of LKP2, and suggests the decoy method can overcome genetic redundancy.

We also measured the effects of ZTL, LKP2, and FKF1 decoy expression on flowering time. We transferred the same plants from the circadian imaging experiment to soil and grew them in inductive long days (16 hour light/8 hour dark). We measured the number of days for the inflorescence to reach 1cm (Fig. 1c, S1b, and S3a), 10 cm (Fig. S3c), or to open the first flower (Fig. S3b). We also measured the number of rosette leaves when inflorescences reached 1cm (Fig. S3d). Plants expressing the FKF1 decoy had the longest delay in flowering (Fig. 1c, S1b and S3 and table S1) (p < .001). Interestingly, LKP2 and ZTL decoys had similar weak effects on flowering time, as both ZTL and LKP2 showed significance in one but not both individual experiments (Fig. 1c, S1b and S3 and table S1). However, when data from both experiments is considered together, the delay in flowering becomes statistically significant for both ZTL and LKP2 decoy plants (p < .001) (Table S1).

Interestingly, some ZTL and LKP2 decoy lines had an approximate 24 hour period but exhibited late flowering (Fig. 1c, S1b, S2c, S3 and S4). We reasoned that these may express the decoy at high levels and sequester GI from the nucleus, as has been shown previously(39, 51). Western blots showing the protein levels of the ZTL decoys confirm this hypothesis, and demonstrate that the late flowering ZTL decoys have the highest decoy expression (Fig. S5). Our results also indicate that the role of ZTL in sequestering GI and causing late flowering may not be dependent on the F-box domain.

### identification of ZTL, LKP2, and FKF1 decoy interacting partners

F-box interactions with substrates are often transient because the substrates dissociate from the F-box or are degraded following ubiquitylation. Therefore, it has been challenging to demonstrate interactions *in vivo* between substrates and the clock F-box proteins, ZTL, LKP2, and FKF1. Recognizing that our decoy system bypasses this challenge and should stabilize interactions in Arabidopsis, we performed immunoprecipitation followed by mass spectrometry (IP-MS) on the ZTL, LKP2, and FKF1 decoy transgenic lines. For all IP-MS experiments, we used the wildtype parental Arabidopsis (Col-0 or Col-0 containing a *CCA1p::Luciferase* construct) and a *35S::His-FLAG-GFP* transgenic line as controls. The wildtype controls for proteins that interact with the column and the *35S::His-FLAG-GFP* transgenic line controls for proteins that interact with the affinity tags.

We first investigated the ZTL decoy because it is well established that ZTL targets the core circadian transcription factor TOC1 for degradation at night(24). We entrained the ZTL decoy in 12 hours light followed by 12 hours dark and collected tissue 6 hours after dark (ZT18) when ZTL, under normal conditions, is actively targeting TOC1 for degradation. We performed IP-MS and detected 439 ZTL peptides, showing we can immunoprecipitate the ZTL decoy bait protein (Table 1). Additionally, we detected 28 TOC1 peptides, showing that the ZTL decoy interacts with a known target. Furthermore, we detected known regulatory partners GI and HSP90(21, 52), suggesting that the decoys are able to form biologically relevant complexes (Table 1).

**Table 1.** Decoy immunoprecipitation followed by mass spectrometry results. IP-MS shows that ZTL, FKF1, and LKP2 decoys interact with known regulatory partners and targets.

| Locus | Protein name | Total spectral counts (from 1 biological replicate) |  |  |
| --- | --- | --- | --- | --- |
|  |  | ZTL decoy | WT (Col-0) | 35S::FLAG-His-GFP |
| Bait |  |  |  |  |
| AT5G57360 | ZTL | 439 | 0 | 0 |
| Regulatory partners |  |  |  |  |
| AT1G22770 | GI | 218 | 0 | 0 |
| AT5G52640 | HSP90.1 | 86 | 120 | 0 |
| AT5G56030 | HSP90.2 | 457 | 278 | 50 |
| AT5G56010 | HSP90.3 | 466 | 0 | 0 |
| AT5G56000 | HSP90.4 | 380 | 240 | 0 |
| Targets |  |  |  |  |
| AT5G61380 | TOC1 | 28 | 0 | 0 |

| Locus | Protein name | Total spectral counts (from 1 biological replicate) |  |  |  |
| --- | --- | --- | --- | --- | --- |
|  |  | LKP2 decoy | FKF1 decoy | WT<br>(CCA1p::LUC+) | 35S::FLAG-His-GFP |
| Bait |  |  |  |  |  |
| AT2G18915 | LKP2 | 1412 | 29 | 0 | 0 |
| AT1G68050 | FKF1 | 0 | 2855 | 128 | 0 |
| Regulatory partners |  |  |  |  |  |
| AT1G22770 | GI | 129 | 265 | 0 | 0 |
| AT5G52640 | HSP90.1 | 51 | 69 | 0 | 0 |
| AT5G56030 | HSP90.2 | 242 | 315 | 0 | 0 |
| AT5G56010 | HSP90.3 | 221 | 314 | 0 | 0 |
| AT5G56000 | HSP90.4 | 192 | 266 | 0 | 0 |
| Targets |  |  |  |  |  |
| AT5G61380 | TOC1 | 74 | 0 | 0 | 0 |
| AT5G24470 | PRR5 | 55 | 0 | 0 | 0 |
| AT5G39660 | CDF2 | 0 | 6 | 0 | 0 |

**Table 1. Decoy immunoprecipitation followed by mass spectrometry results (continued)**
| Locus | Protein name | Total spectral counts (from 2 biological replicates) |  |  |  |  |
| --- | --- | --- | --- | --- | --- | --- |
|  |  | ZTL decoy |  |  | WT<br>(CCA1p::LUC+) | 35S::FLAG-His-<br>GFP |
|  |  | 3h before dusk<br>(LD) | 3h after dusk<br>(LD) | 3h after subjective<br>dusk (LL) |  |  |
| Bait |  |  |  |  |  |  |
| AT5G57360 | ZTL | 218 | 298 | 190 | 0 | 0 |
| Regulatory partners |  |  |  |  |  |  |
| AT1G22770 | GI | 51 | 75 | 34 | 0 | 0 |
| AT1G22770 | HSP90.1 | 59 | 44 | 47 | 21 | 6 |
| AT5G52640 | HSP90.2 | 144 | 172 | 122 | 185 | 88 |
| AT5G56030 | HSP90.3 | 83 | 98 | 113 | 35 | 0 |
| AT5G56010 | HSP90.4 | 108 | 79 | 95 | 0 | 103 |
| Targets |  |  |  |  |  |  |
| AT5G08330 | CHE | 0 | 19 | 17 | 10 | 0 |

We next examined the interacting proteins of LKP2 and FKF1 decoys via IP-MS. Because LKP2 and FKF1 are predicted to regulate seasonal flowering time through degradation of CDF proteins(14), we grew these lines in inductive long-day conditions and collected samples two hours after dark (ZT18). We performed IP-MS and detected 1412 LKP2 and 2855 FKF1 peptides, demonstrating immunoprecipitation of the LKP2 and FKF1 decoy bait proteins (Table 1). Again we were able to detect GI and HSP90(20), two known regulatory partners, suggesting the decoys are forming biologically relevant protein complexes.

In the LKP2 decoy IP, we detected 74 TOC1 and 55 PRR5 peptides, suggesting that LKP2 can interact with clock transcription factors in Arabidopsis. The FKF1 decoy IP failed to detect clock proteins but did identify 6 peptides from CDF2, a known FKF1 target that regulates seasonal flowering time (Table 1). These results correlate with the genetic experiments showing that LKP2 has a strong effect on circadian clock function and FKF1 has a strong effect on flowering. While we were successful at identifying some high confidence targets of ZTL, FKF1, and LKP2, we were not able to identify all known interactors(31, 39, 40, 53-59) (Table S2). This is likely due to the age of the tissue, collection times, or tissue-type that we used to perform the IP-MS.

We next asked whether we could detect ZTL target proteins that have not been identified previously. To do this, we performed a more comprehensive time course IP-MS experiment. We entrained the ZTL decoy in 12 hour light/12 hour dark growth conditions. We collected tissue in the light (three hours before dusk, ZT9) and dark (three hours after dusk, ZT15), then transferred the plants to constant light for 24 hours and collected tissue in the subjective dark (three hours after subjective dusk, ZT15 in LL conditions). We detected its known regulatory partners GI and HSP90 at all time points (Table 1). To detect ZTL substrates that have not been previously studied, we filtered our interaction list for transcription factors or transcriptional regulators involved in the circadian clock(46, 60-69) (Tables 1 and S3). We found multiple transcription factors of interest including a known TOC1 interaction partner, CCA1 HIKING EXPEDITION (CHE)(46). Of note, CHE only immunoprecipitated with ZTL at late time points (three hours after dusk and three hours after subjective dusk), consistent with what we would expect for a ZTL degradational target. As CHE is enriched in the ZTL IPs (Table 1), regulates *CCA1* expression, and is known to interact with TOC1(46), we performed follow-up experiments to determine if it is a bona fide ZTL target.

### Use of heterologous systems to validate interaction and ubiquitylation

As substrate proteins often interact directly with the substrate recognition domain of an E3 ligase, we tested whether CHE interacts directly with ZTL using yeast two-hybrid assays. In yeast, CHE interacts directly with the full length and decoy ZTL (Fig. 2a). Moreover, CHE interacts with the LOV domain of ZTL, similarly to TOC1(24). TOC1 is a light-dependent target of ZTL, making it likely that CHE is ubiquitylated and degraded in a light-dependent manner similarly to TOC1 (Fig. 2a).

**Figure 2.**
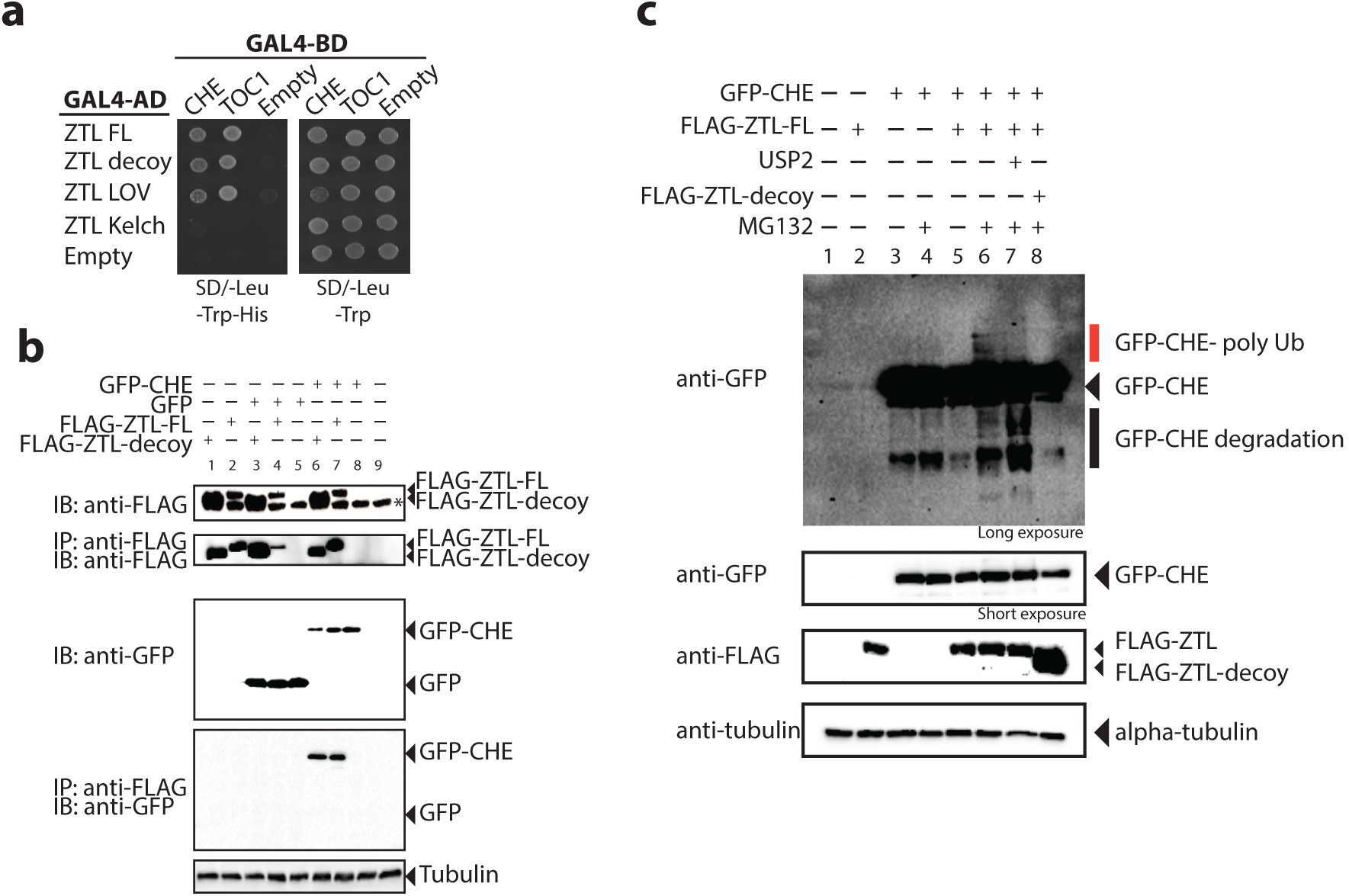
ZTL interacts directly with CHE and mediates its ubiquitylation. (a) Yeast two-hybrid assays between ZTL and CHE or TOC1. FL= Full length, decoy= translationally fused LOV and Kelch-repeat domains, LOV= LOV domain only, Kelch= Kelch-repeat domain only. (b) Co-immunoprecipitation experiments between CHE and ZTL performed in HEK293T cells. GFP-CHE, or GFP alone, was coexpressed with full-length or decoy ZTL translationally fused to a FLAG affinity tag. Immunoprecipitation was performed with the FLAG tag. IP=immunoprecipitation, IB=immunoblot. The star (*) present in the FLAG-input blot marks a nonspecific band. (c) Ubiquitylation assay performed in mammalian tissue culture cells. GFP-CHE was co-expressed with full-length or decoy ZTL. They were expressed in the presence or absence of 30 μΜ MG132 or 2 μg USP2cc. Red line denotes poly-ubiquitylated forms of CHE. FL=Full length.

Ubiquitylation studies are often technically difficult and time-consuming to perform and interpret in plant model systems. In order to determine whether ZTL mediates ubiquitylation of CHE, we used mammalian tissue culture cells as a heterologous ubiquitylation system. Mammalian tissue culture has three distinct advantages for testing the relationship between plant F-box proteins and potential targets: 1) the mammalian UPS supports ubiquitylation by plant F-box proteins, 2) redundant plant proteins do not complicate the interpretation of results, and 3) there are a wealth of reagents available for protein expression and ubiquitylation assays in mammalian cells.

We first used this system to test whether CHE interacts with ZTL in mammalian cells. We co-expressed CHE and either full length or decoy ZTL in HEK 293T cells. Figure 2b shows that FLAG-tagged full length ZTL, FLAG-tagged ZTL decoy, and GFP-tagged CHE can be expressed in HEK293T cells. We immunoprecipitated ZTL using an anti-FLAG antibody and then performed an immunoblot with anti-GFP to determine if CHE interacts with ZTL. We detected CHE interaction with the full-length and decoy ZTL (Fig. 2b lanes 6-7), but this interaction was not present in the GFP only control (Fig. 2b lanes 3-5) or when CHE-GFP is expressed alone (Fig. 2b lane 8). This supports the hypothesis that CHE can directly interact with ZTL and allows us to test whether ZTL mediates ubiquitylation of CHE in mammalian cells.

To determine whether ZTL mediates ubiquitylation of CHE, we co-expressed GFP-CHE and FLAG-ZTL in the presence or absence of the proteasome inhibitor MG132 (Fig. 2c). When full length ZTL is coexpressed with CHE in the presence of a proteasome inhibitor, a higher molecular weight laddering of the CHE protein occurs (Fig. 2c lane 6). This laddering diminishes upon addition of a general deubiquitylating enzyme, USP2cc, to the lysate (Fig. 2c lane 7), demonstrating that these bands are ubiquitylated forms of CHE. Furthermore, the ZTL-dependent laddering of CHE is blocked by the ZTL decoy (Fig. 2c lane 8) demonstrating that our decoy inhibits the function of the full-length protein as predicted.

### *In planta* study of identified F-box targets

Based on our interaction and ubiquitylation studies, we hypothesize that CHE protein will be destabilized in the dark when ZTL is actively degrading target proteins(24, 29). This would result in cycling of CHE protein levels in light/dark conditions even in the absence of transcriptional changes. To test this we constitutively expressed *CHE-GFP* under a *35S* promoter (*35S::CHE:GFP*) in Arabidopsis (Fig. 3a) and performed time course western blotting under three different light regimes: 12 hour light/12 hour dark (LD), constant light (LL), and constant dark (DD) (Fig. 3b-c). In LD conditions, CHE protein levels cycle robustly, peaking in the light (Fig. 3b). In constant light, CHE protein levels do not show robust 24 hour cycles but remain at a constant high level (Fig. 3c), and in constant dark, CHE protein levels are undetectable at all time points (Fig. 3d). In order to cross-compare protein levels from the three time course experiments, we loaded the sample from 9 hours after lights on (light/dark) or subjective lights on (constant light and constant dark) in the western blots of the other two time courses (Fig. 3b-d lanes labeled LL, LD, or DD). This demonstrates that CHE protein levels are controlled by light cycles, which control ZTL activity. This suggests that although CHE mRNA expression is controlled by the circadian clock, CHE protein stability is controlled by light/dark cycles.

**Figure 3.**
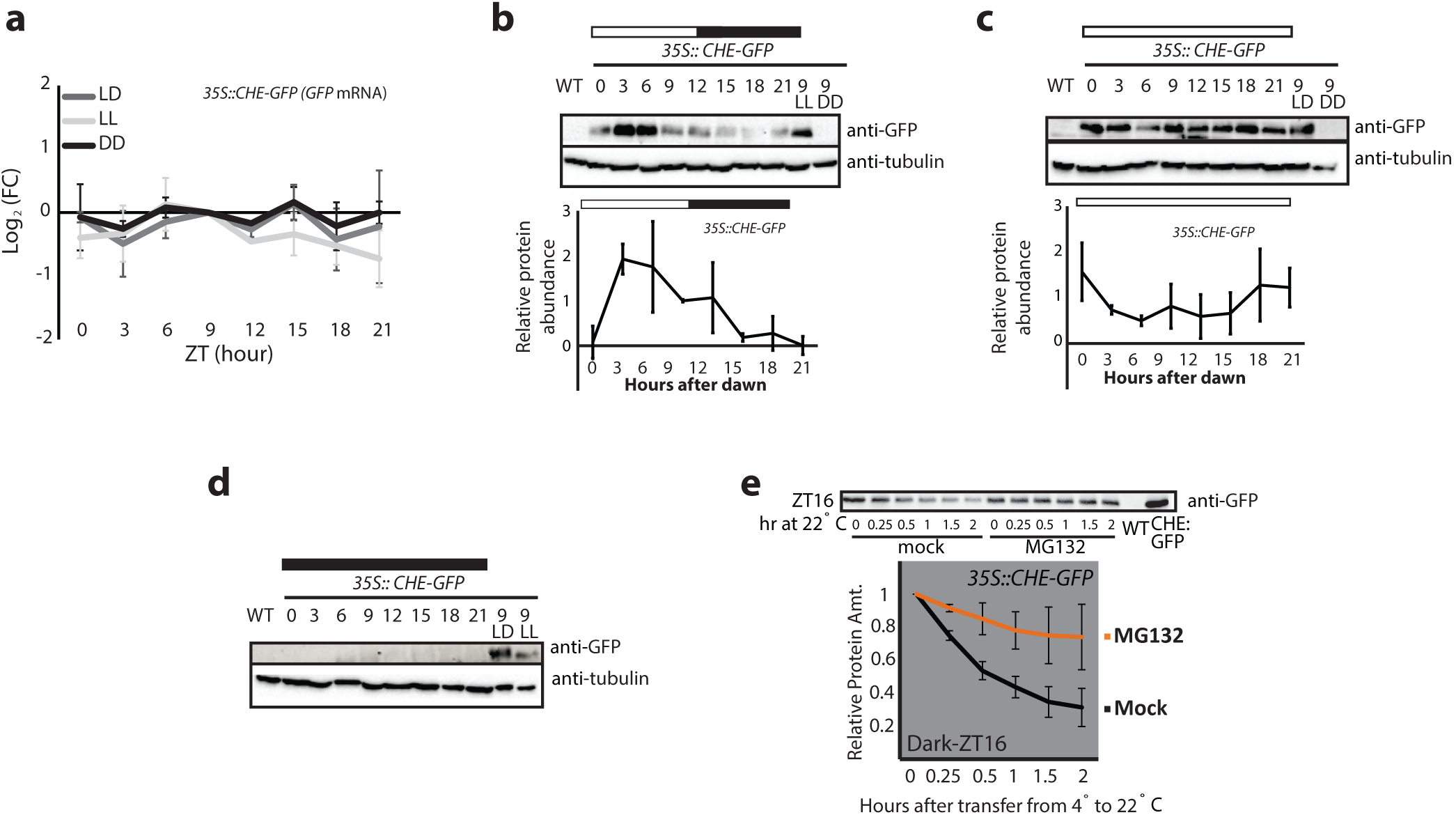
CHE protein cycles over daily time courses. (a) *CHE* mRNA expression was measured using quantitative RT-PCR in a *35S::CHE-GFP* transgenic line in LD (12 hour light/12 hour dark), LL (constant light), and DD (constant dark) growth conditions. (b-d) CHE protein levels were measured using western blotting in a *35S::CHE-GFP* transgenic line in (b) LD (12 hour light/12 hour dark), (c) LL (constant light), and (d) DD (constant dark) growth conditions. Quantifications are the average of three biological replicates with error bars showing standard deviation. DD levels were too low for quantification. ZT9 samples are presented for cross-comparison of relative levels of protein between LD, LL, and DD time courses. (e) Cell-free degradation was performed on *35S::CHE-GFP* grown in LD cycles. Sample was collected at ZT16 in the dark and protein was extracted at 4^o^C and split into tubes at room temperature for mock treatment or treatment with 1μM MG132 for the indicated times.

It is possible that CHE is stabilized in the light rather than destabilized in the dark. To determine whether CHE is degraded by the proteasome in the dark, we performed a cell-free degradation assay on *35S::CHE-GFP* tissue collected 4 hours after dusk (ZT16) in a 12 hour light/12 hour dark time course. We separated the tissue into a mock and MG132-containing sample buffer and measured protein stability at various time points over two hours via western blotting (Fig. 3e). MG132 stabilizes CHE-GFP, demonstrating that the ubiquitin proteasome system is acting on CHE stability at night during ZTL’s peak activity. This has been demonstrated from known ZTL targets, such as TOC1(24).

## Discussion

The decoy method can rapidly determine the genetic contribution of any F-box protein. Three challenges inhibit progress in studying plant F-box proteins: genetic redundancy, transient interactions with substrate proteins, and validation of substrates. The decoy method presented here inverts the function of an E3 ubiquitin ligase, stabilizing substrate proteins even in the presence of the endogenous F-box or functionally redundant F-box proteins. This creates a dominant-negative genetic effect that can be used to determine the functions of any F-box protein family, even in the presence of functional redundancy. This dominant-negative feature of our technique will be especially useful in plant species and other species with genome duplications, where redundancy is prevalent. Furthermore, this platform presents a means for rapid identification of F-box substrate proteins using IP-MS and a subsequent method for validation of substrate ubiquitylation by the assayed F-box protein.

The decoy method reveals the hidden genetics of LOV/F-box/Kelch-repeat proteins in plants. ZTL, LKP2, and FKF1 have a complicated genetic relationship(15, 40). Our analysis using decoys has confirmed the roles of ZTL and FKF1 in the circadian clock and flowering time, and has also shown that LKP2 plays an important role in regulating clock period. This provides further proof that ZTL, LKP2, and FKF1 have diverged into specialized roles in the circadian clock and flowering. LKP2 isn’t fully redundant with ZTL or FKF1 in Arabidopsis, but, intriguingly, *Brassica rapa* has three copies of LKP2 and no ZTL or FKF1(70), suggesting that LKP2 is sufficient to drive seasonal flowering and maintain circadian period in other plant species. It is reasonable to believe that complicated genetic scenarios such as these are prevalent across F-box families in all plant species. Beyond traditional knockout and overexpression studies, decoys should become an additional method to unravel complicated genetic and biochemical structures in protein degradation families.

Our results suggest that LKP2 has the potential to play a significant role in regulating clock period length. Expression of the LKP2 decoy shifts the clock period by nearly two hours, an effect not seen in knockout or full-length overexpression studies (Fig. 1b and S1). Multiple lines expressing the decoys at various levels have the same clock effect, thus one can hypothesize that post-translational mechanisms account for the difference in period between ZTL and LKP2 decoys. It will be interesting to investigate this idea, perhaps in a heterologous system such as with the mammalian tissue culture approach described here. The detection of PRR5 and TOC1 in the LKP2 decoy IP-MS does suggest that LKP2 can target clock factors *in planta.* We also detect endogenous LKP2 interacting with the ZTL decoy in IP-MS, showing that the endogenous protein is expressed despite predicted low relative mRNA expression.

The decoy method can reveal unknown targets for F-box proteins. F-box proteins can target multiple substrates and substrate types, but it is hard to detect F-box targets *in vivo* in an unbiased manner. PRR5 and TOC1 are well known homologous transcription factor targets of ZTL that regulate the circadian clock, and the CDFs are well known targets of FKF1 that regulate seasonal flowering. Our decoy approach revealed that CHE is a previously unknown target of ZTL. CHE is a TCP-type transcription factor that interacts with TOC1 to control *CCA1* expression, but has no apparent sequence similarity with the PRRs. This suggests that ZTL can regulate diverse targets and is likely a master regulator of nighttime repressors of *CCA1.*

The results presented here strongly suggest that CHE is a bona fide ZTL target. Although outside of the scope of this manuscript, further validation of CHE as a degradational substrate of ZTL is planned. Measuring the daily stability of CHE protein in both *ztl* and *gi* mutant backgrounds would provide genetic evidence that ZTL degrades CHE in a light-dependent manner *in vivo* in a fashion analogous to the previously known ZTL substrates, TOC1 and PRR5. As with other oscillatory biological processes, E3 ubiquitin ligases with multiple substrates can target them sequentially. It will be interesting to use our *in vivo* and heterologous expression systems to test the order of degradation of ZTL targets during the day.

Furthermore, as with other ZTL substrates, it will be important to determine whether CHE can be ubiquitylated by LKP2 or FKF1.

The interaction with LUX ARRHYTHMO (LUX, also known as PCL1) and BROTHER OF LUX ARRHYTHMO (BOA) (Table S3), core components of the circadian clock evening complex(66), is intriguing and suggests that further investigation into the targets of ZTL is imperative to understand its complete role in clock function. It is also possible that in our list of ZTL, LKP2, and FKF1 interacting proteins (Tables 1 and S2-S6) there may be new clock or flowering time regulators that haven’t been identified by other means. Surprisingly, we detected interaction between ZTL, LKP2, and FKF1 in our studies. This has been controversial in the past(33, 39, 53), but our work suggests they may form higher order complexes *in vivo* possibly to regulate target stability or regulate their own stability. Future studies will likely tease apart the functional significance of these putative dimerization events, and the decoy strategy provides a unique opportunity to study this *in vivo.*

Heterologous ubiquitylation assays complement the decoy method as a rapid target validation approach. We demonstrated the ability to reconstitute CHE ubiquitylation by ZTL in HEK293T cells, a mammalian tissue culture system (Fig. 2b and 2c). One of the drawbacks of IP-MS approaches is the prevalence of false positives that cloud interpretation of results. Mammalian tissue culture is a rapid eukaryotic protein expression system that can be used to demonstrate the ubiquitylation of a substrate by a plant F-box protein. This allows for rapid screening of potential F-box targets to quickly generate hypotheses that can then be tested in plant systems (as in Fig. 3), which are traditionally time-consuming. This method also reduces the possibility of indirect interactions through interacting partners of the F-box protein or target of interest which would be present in an *in vivo* assay. The combined decoy and heterologous ubiquitylation assays is a powerful combinatorial method that can be used to understand the role of any E3 ubiquitin ligase from plants.

The decoy method can be used for high-throughput screening of F-box protein function. The results presented here demonstrate the effectiveness of using the decoy approach to break apart redundancy within a gene family. There are additional benefits of this approach that make it amenable as a screening platform. First, nearly all F-box proteins have the F-box domain in the N-terminus, providing a simple two primer PCR strategy for high-throughput cloning of F-box decoys. Second, a screen can be done in the T1 generation because of the dominant genetic effects of decoys (Fig. 1b and 1c). Third, T1 lines of varying expression can be assayed for phenotypes, reducing the danger of expression artifacts. Finally, genetic screening can be followed by rapid IP-MS experiments to determine the interacting proteins that may be stabilized by the decoy F-box. We used overexpression with the *35S* promoter because ZTL, FKF1, LKP2 are likely to function broadly, and the strong *35S* promoter can effectively overcome any redundancy to reveal all possible functions of this family. Additionally, an EMS mutagenesis allele in the ZTL F-box domain has been identified previously(71). This allele is dominant-negative and leads to a 28 hour period phenotype similar to the ZTL decoy, suggesting constitutive expression is not causing aberrant phenotypes. Sometimes a strong ubiquitous promoter might be problematic, such as for proteins with a highly restricted expression domain; in these cases our system is easily adapted to use the native promoter to drive expression.

IP-MS is a streamlined follow-up approach to identify substrates of E3 ligases. Previous approaches have used point mutations in the domain that recruits the ubiquitylation machinery followed by IP-MS to detect target interactions(50). The advantage of this strategy is that the domain may play additional roles in interacting with substrate proteins or other regulatory partners. The drawback of this approach is that point mutations may not fully prevent interaction with the rest of the SCF complex, allowing for residual degradation of substrates. Additionally, it is not feasible for high-throughput screening approaches. The decoy strategy is amenable to high throughput strategies, and our results suggest it is effective at revealing the interacting partners and genetic functions of E3 ubiquitin ligases.

Decoys can be employed in any biological system, but may have greater impact on traditionally intractable systems such as polyploid plants that are plagued by genetic redundancy. Some of the most important food sources, such as hexaploid wheat, are traditionally difficult to manipulate genetically. Dominant-negative approaches, such as decoys, may be used to great effect to improve high value characteristics. The ability to rapidly identify the functions of the F-box proteins through IP-MS is also now possible in those species for which we have genome sequences.

---

**Figure S1.**
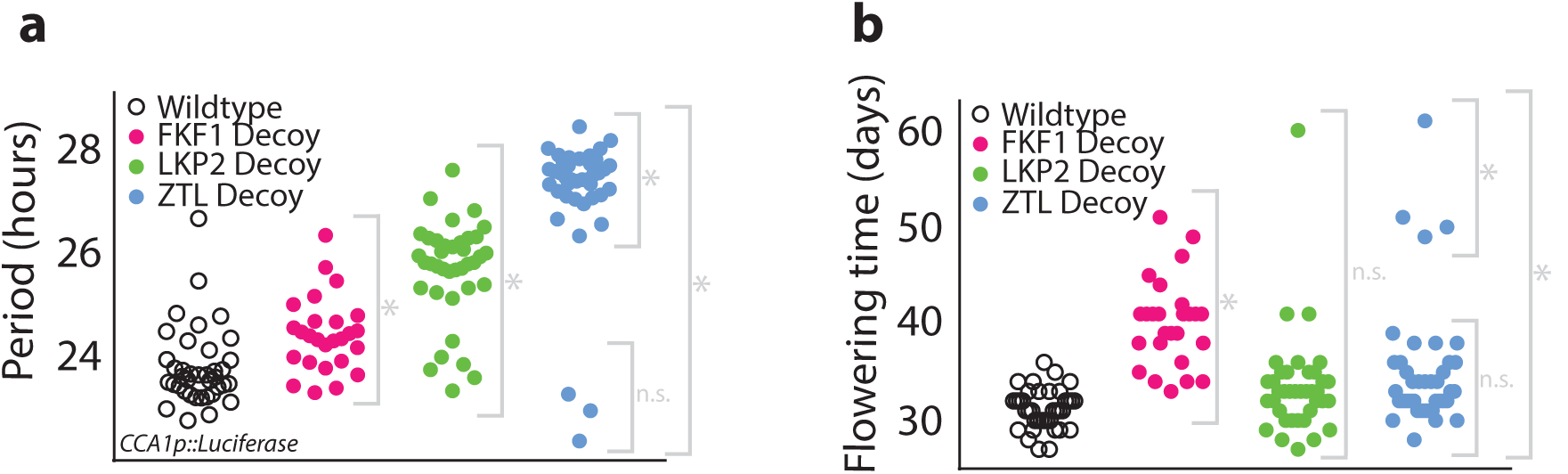
Decoys reveal the genetic contributions of redundant F-box genes. Experiments were performed as in Figure 1 with different T1 plants. (a) Period lengths (as measured by *CCA1p::Luciferase* activity) for T1 transgenic decoys from ZTL (blue), LKP2 (green), FKF1 (pink), and the wildtype parental line (*CCA1p::Luciferase* - open circles). (b) Flowering time (number of days for inflorescence stem to reach 1cm) for T1 transgenic decoys from ZTL (blue), LKP2 (green), FKF1 (pink), and the wildtype parental line (*CCA1p::Luciferase-open* circles). Grey brackets define individual groups used for statistical testing against the wildtype control using a Welch’s t-test with a Bonferroni-corrected α of 1.67×10^−3^. * represents p < α, n.s p > α. HF = His-FLAG. F = F-box.

**Figure S2.**
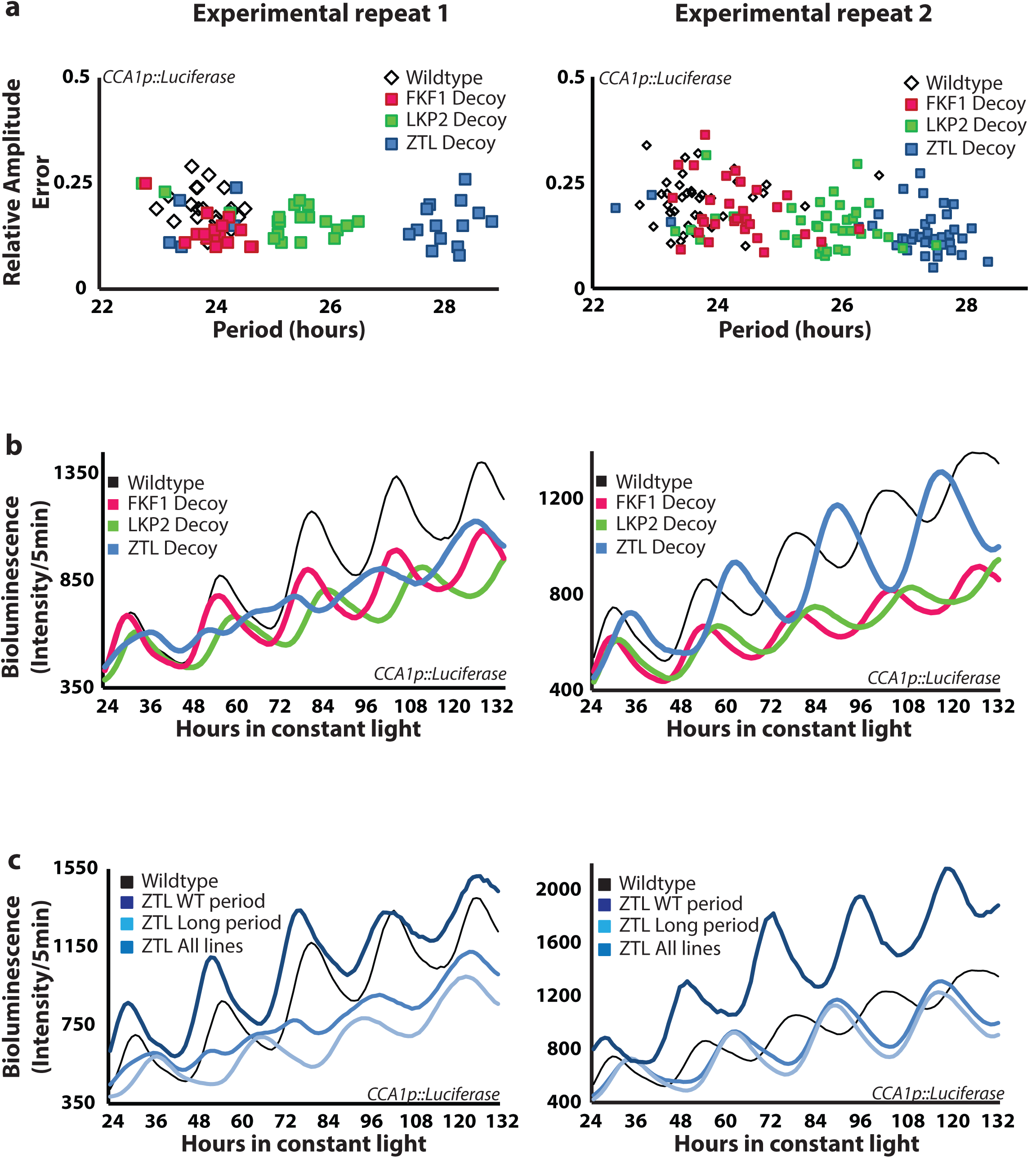
Decoys reveal the genetic contributions of redundant F-box genes to the circadian clock. (a) Period data from figures 1 (repeat 1) and S1 (repeat 2) plotted against relative amplitude demonstrates that all lines tested are considered rhythmic (relative amplitude error less than 0.5). (b) Average traces from wildtype (parental *CCA1p::Luciferase)* and the ZTL, LKP2, and FKF1 decoys from figures 1 and S1. (c) Average traces from wildtype (parental *CCA1p::Luciferase),* all ZTL decoy lines, long period ZTL decoy lines, and wildtype period ZTL decoy lines.

**Figure S3.**
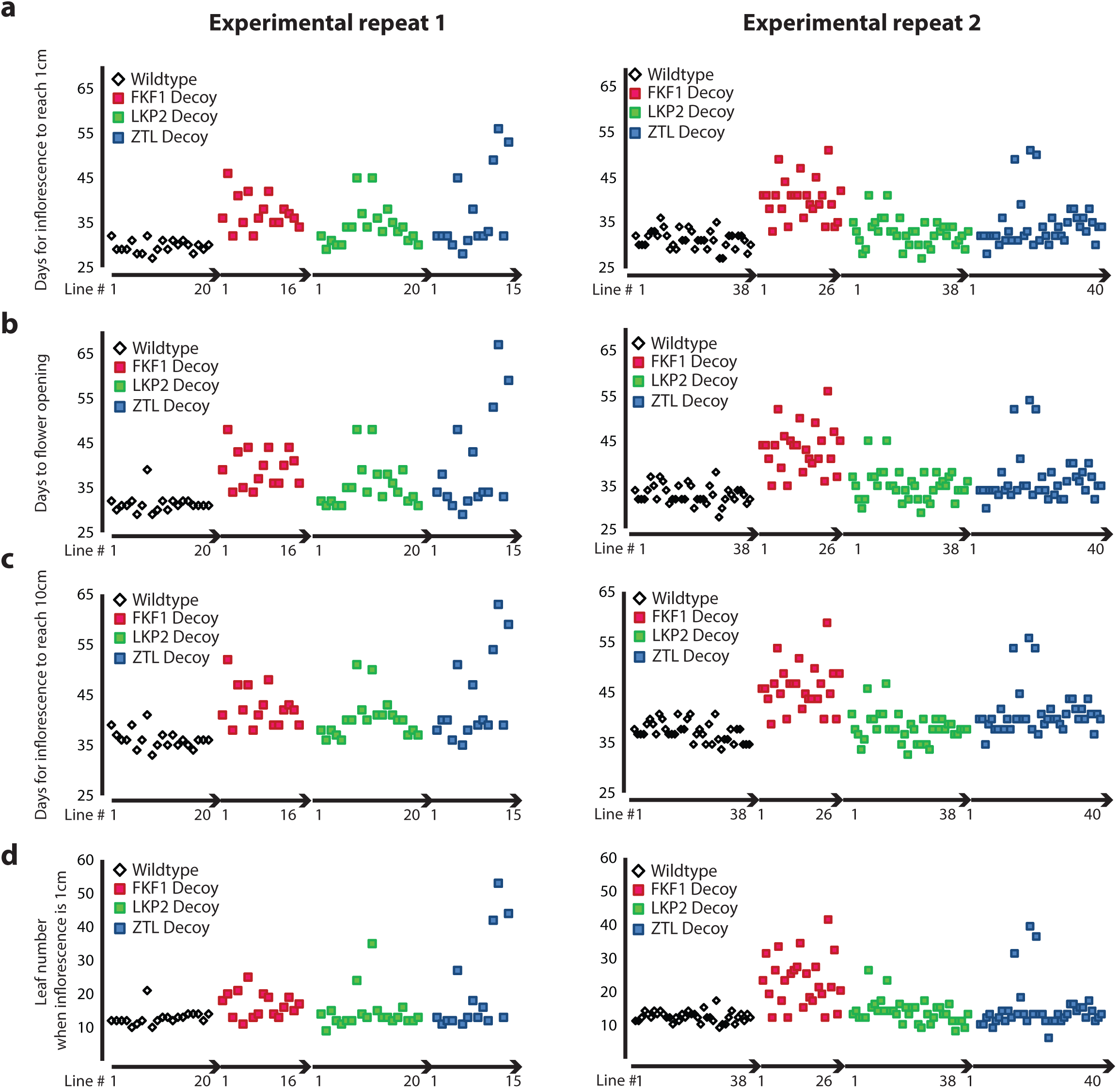
Decoys reveal the genetic contributions of redundant F-box genes to seasonal flowering time. Flowering time was measured for wildtype and the ZTL, LKP2, and FKF1 decoys using four parameters: (a) number of days for the inflorescence stem to reach 1 cm, (b) number of days to the first flower opening, (c) number of days for the inflorescence stem to reach 10 cm, and (d) number of rosette leaves when the inflorescence stem reaches 1 cm. Individual T1 lines are plotted on the x-axis so lines can be compared across all parameters.

**Figure S4.**
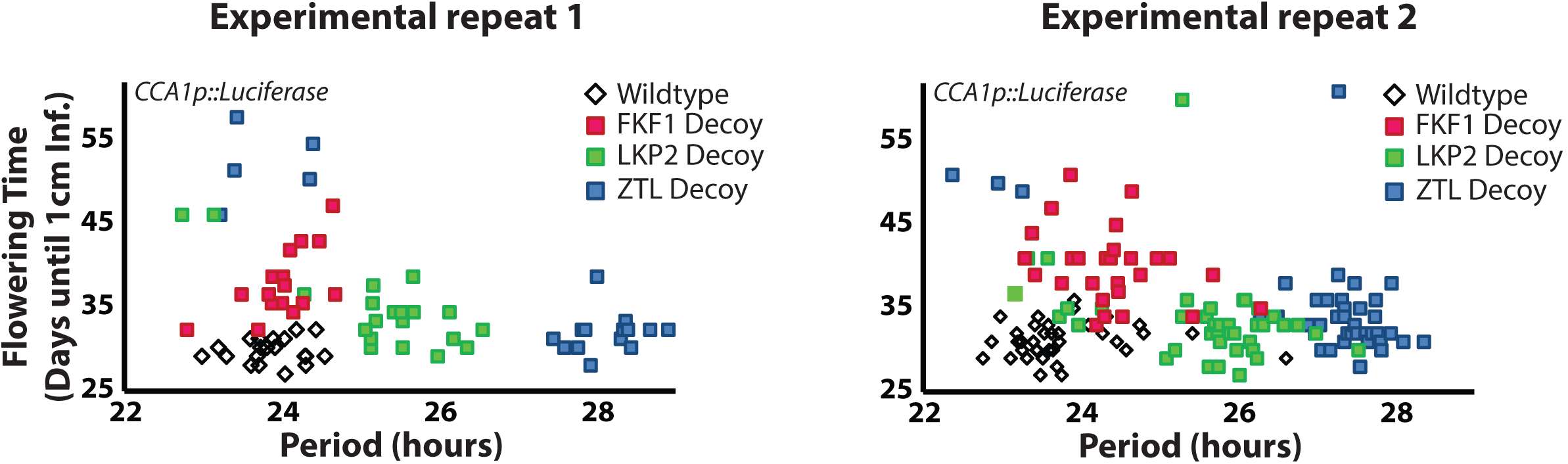
Decoys reveal the genetic contributions of redundant F-box genes to the clock and seasonal flowering time. Period length was plotted against flowering time for experiments shown in figures 1 (b and c) and S1 (a and b).

**Figure S5.**
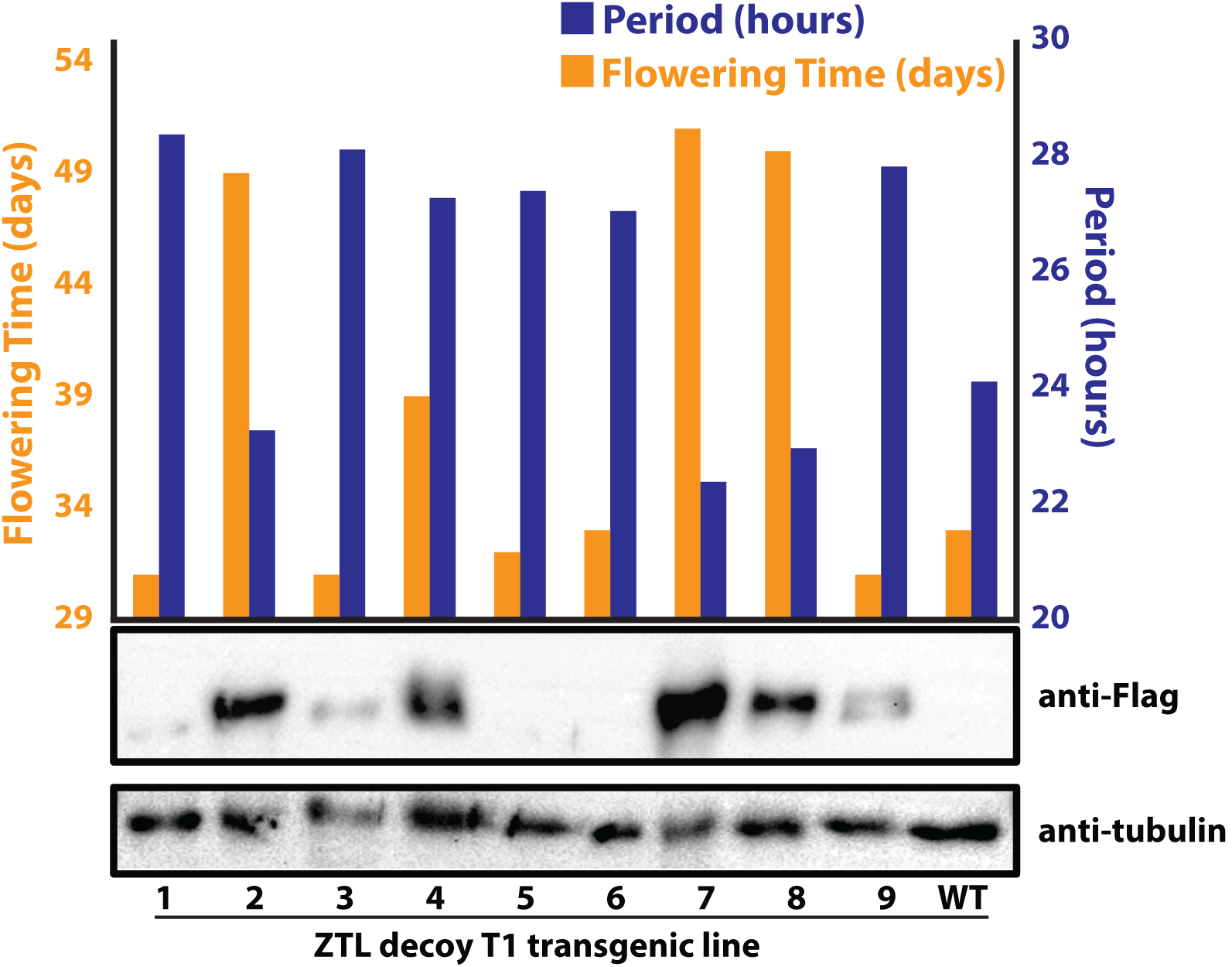
Very late flowering ZTL decoy lines express the decoy at high levels. (Top) Flowering time and period are plotted for 10 individual ZTL decoy lines. 3 lines (#2,7,9) have wildtype period but flower late. (Bottom) Western blot showing that lines #2,7, and 9 express the ZTL decoy at high levels. *WT=CCA1p::Luciferase* parental line.

**Table S1.** Genetics Experiments Statistics. Statistical analysis of the period effects and flowering time effects of the ZTL, LKP2, and FKF1 decoy lines from figures 1b, 1c, S1a and S1b. Null hypothesis: the decoy population has the same period or flowering time as wildtype. p-values are from an unpaired two-tailed Welch’s t-test with a Bonferroni-corrected α of 1.67×10^−3^.

| Experimental Repeat 1 |  | N | Circadian |  | Flowering (1cm inf.) |  |
| --- | --- | --- | --- | --- | --- | --- |
| Genotype | Group |  | Mean Period (hrs) | P-Value | Mean (days) | P-Value |
| ZTL decoy | All | 19 | 26.96 | $2.46 \times 10^{-6}$ | 36.6 | $3.45 \times 10^{-3}$ |
| | Long period | 14 | 28.11 | $2.18 \times 10^{-22}$ | 31.6 | $7.83 \times 10^{-3}$ (n.s.) |
| | WT period | 5 | 23.77 | $7.12 \times 10^{-1}$ (n.s.) | 50.6 | $2.77 \times 10^{-4}$ |
| FKF1 decoy | All | 16 | 24.02 | $5.07 \times 10^{-2}$ (n.s.) | 37.2 | $6.36 \times 10^{-7}$ |
| LKP2 decoy | All | 20 | 25.27 | $2.29 \times 10^{-6}$ | 34.2 | $2.49 \times 10^{-4}$ |
| WT (*CCA1p::Luc+*) | All | 20 | 23.87 | -- | 29.7 | -- |

| Experimental Repeat 2 |  | N | Circadian |  | Flowering (1cm inf.) |  |
| --- | --- | --- | --- | --- | --- | --- |
| Genotype | Group |  | Mean Period (hrs) | P-Value | Mean (days) | P-Value |
| ZTL decoy | All | 40 | 27.07 | $5.93 \times 10^{-21}$ | 35.2 | $6.33 \times 10^{-4}$ |
| | Long period | 37 | 27.41 | $1.53 \times 10^{-34}$ | 34 | $3.80 \times 10^{-3}$ |
| | WT period | 3 | 22.86 | $5.31 \times 10^{-2}$ (n.s.) | 50 | $2.64 \times 10^{-5}$ |
| FKF1 decoy | All | 27 | 24.40 | $8.11 \times 10^{-4}$ | 40 | $4.50 \times 10^{-11}$ |
| LKP2 decoy | All | 38 | 25.63 | $2.62 \times 10^{-14}$ | 33.5 | $1.64 \times 10^{-2}$ (n.s.) |
| WT (*CCA1p::Luc+*) | All | 38 | 23.76 | -- | 31.2 | -- |

| Experiments 1 and 2 Combined |  | N | Circadian |  | Flowering (1cm inf.) |  |
| --- | --- | --- | --- | --- | --- | --- |
| Genotype | Group |  | Mean Period (hrs) | P-Value | Mean (days) | P-Value |
| ZTL decoy | All | 59 | 27.03 | $1.97 \times 10^{-24}$ | 35.66 | $4.30 \times 10^{-6}$ |
| | Long period | 51 | 27.60 | $9.17 \times 10^{-59}$ | 33.35 | $3.17 \times 10^{-4}$ |
| | WT period | 8 | 23.43 | $1.76 \times 10^{-1}$ (n.s.) | 50.38 | $2.06 \times 10^{-7}$ |
| FKF1 decoy | All | 43 | 24.26 | $6.46 \times 10^{-4}$ | 38.95 | $4.17 \times 10^{-16}$ |
| LKP2 decoy | All | 58 | 25.50 | $1.82 \times 10^{-19}$ | 33.72 | $4.67 \times 10^{-5}$ |
| WT (*CCA1p::Luc+*) | All | 58 | 23.69 | -- | 30.66 | -- |

**Table S2.**
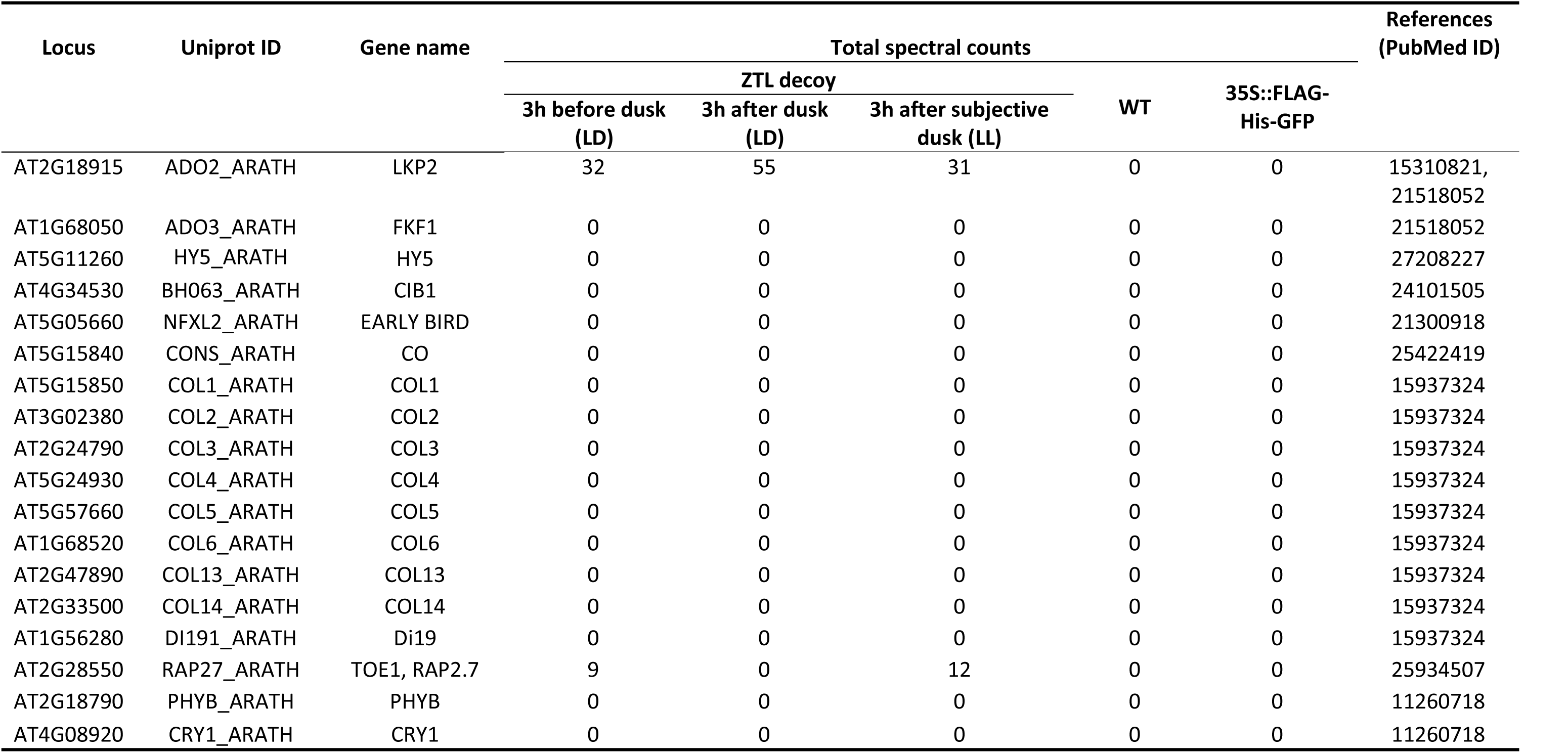
Published interactors in ZTL decoy IP-MS. Spectral counts of published ZTL interactors in time course ZTL decoy IP-MS experiments.

**Table S3.** Transcription factors and transcription regulators involved in clock function that co-immunoprecipitated with ZTL decoy. Transcription factors identified in ZTL decoy IP-MS experiments that are known to be involved in clock function or response to light.

| Locus | Uniprot ID | Gene name | Protein family | Total spectral counts |  |  |  |  |
| --- | --- | --- | --- | --- | --- | --- | --- | --- |
|  |  |  |  | ZTL decoy |  |  | WT | 35S::FLAG<br>-His-GFP |
|  |  |  |  | 3h before<br>dusk (LD) | 3h after<br>dusk (LD) | 3h after subjective<br>dusk (LL) |  |  |
| Transcription factor |  |  |  |  |  |  |  |  |
| AT1G35460 | BH080_ARATH | FLOWERING BHLH 1 (FBH1) | bHLH | 34 | 22 | 34 | 19 | 0 |
| AT1G72010 | TCP22_ARATH | TCP22 | TCP | 15 | 0 | 64 | 27 | 8 |
| AT5G08330 | TCP21_ARATH | CCA1 HIKING EXPEDITION (CHE) | TCP | 0 | 19 | 17 | 10 | 0 |
| AT3G46640 | PCL1_ARATH | LUX, PHYTOCLOCK 1 (PCL1) | MYB,<br>G2-like | 20 | 0 | 0 | 0 | 0 |
| AT5G59570 | PCLL_ARATH | BROTHER OF LUX ARRHYTHMO (BOA) | MYB,<br>G2-like | 0 | 20 | 0 | 0 | 0 |
| Transcriptional regulator |  |  |  |  |  |  |  |  |
| AT1G12910 | LWD1_ARATH | ANTHOCYANIN11 (ATAN11) | WD40 | 42 | 55 | 46 | 19 | 0 |
| AT3G26640 | LWD2_ARATH | LIGHT-REGULATED WD 2 (LWD2) | WD40 | 26 | 37 | 34 | 0 | 0 |

**Table S4** IP-MS data for the ZTL decoy at six hours after dusk in 12 hour light/12 hour dark conditions (ZT18) with controls.

**Table S5** IP-MS data for the LKP2 and FKF1 decoys at two hours after dusk in long day conditions (ZT18 in 16 hour light/8 hour dark) with controls.

**Table S6** IP-MS data for the ZTL decoy at three hours before dusk (ZT9 in LD), three hours after dusk (ZT15 in LD), and three hours after subjective dusk (ZT 15 in LL) with controls.

**Table S7.**
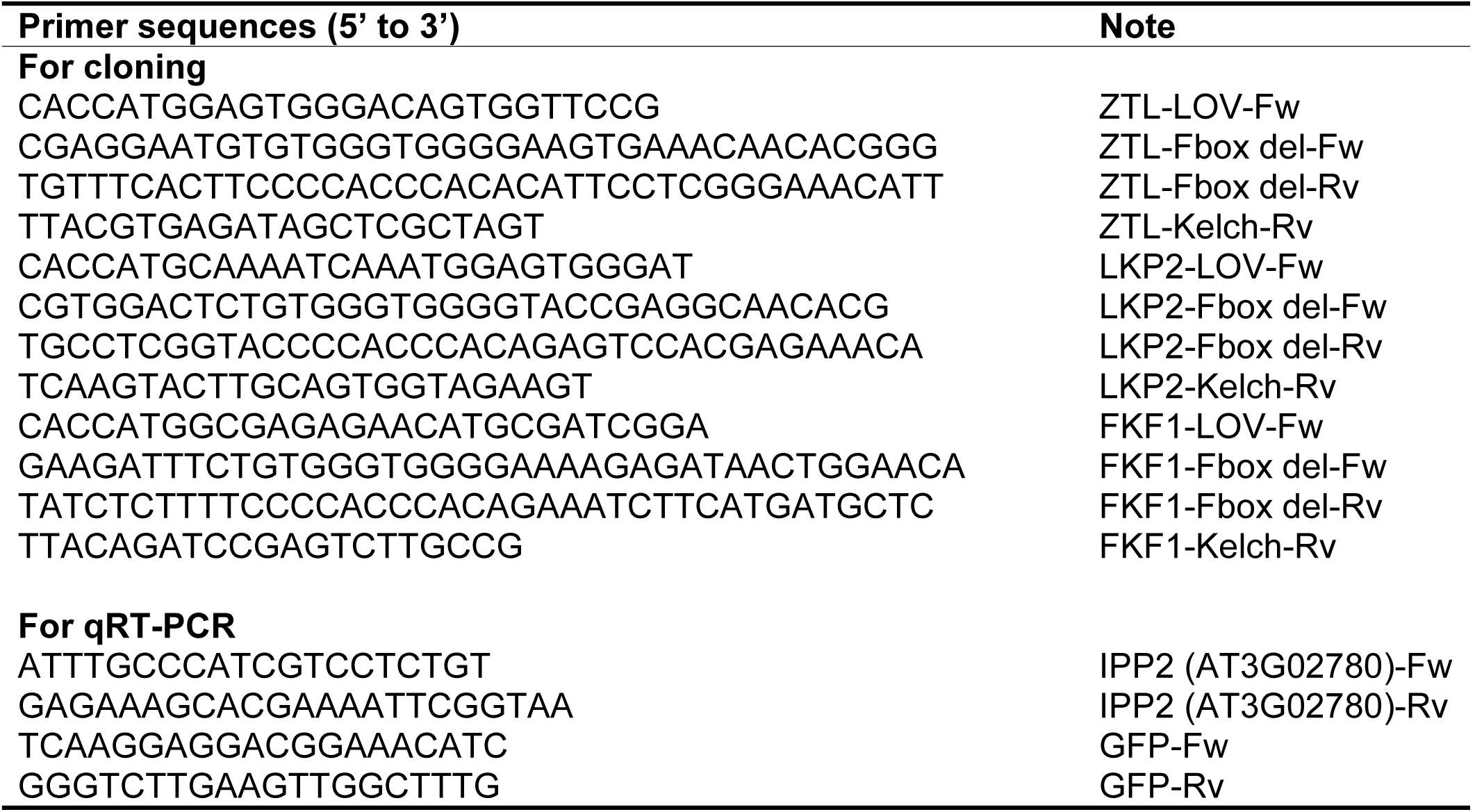
Primer sequences. Primers used for cloning decoy constructs and quantitative RT-PCR experiments.

## Methods

### Plant materials and growth conditions

The decoy constructs for ZTL (AT5G57360), LKP2 (AT2G18915), and FKF1 (AT1G68050) were constructed by fusion of the LOV and Kelch domains using overlap extension PCR with the primers listed in Table S7. The design, including amino acid numbers for ZTL, LKP2, and FKF1 decoys, are shown in Figure 1a. The PCR products were cloned into pENTR/D-TOPO vectors (Invitrogen, cat. # K240020). The decoys were then fused to FLAG and His tags at the N-terminus and under the control of a CaMV 35S promoter by recombination into the plant binary pDEST vector pB7-HFN(72) using LR recombination. The decoy constructs were transformed into Arabidopsis Col-0 expressing the circadian reporter *CCA1p::Luciferase* (46) or Col-0 by the floral-dip method(73) using *Agrobacterium tumefaciens* GV3101. The *35S::CHE-GFP* transgenic lines were generated previously(46).

For growth of Arabidopsis seedlings, Arabidopsis seeds were surface sterilized in 70% ethanol and 0.01% Triton X-100, sown on ½ MS plates (2.15 g/L Murashige and Skoog medium, pH 5.7, Caisson Laboratories, cat# MSP01 and 0.8% bacteriological agar, AmericanBio, cat# AB01185) and stratified at 4 °C for 2 days. Following stratification, seeds were grown at 22 °C in 12 hour light/12 hour dark cycles at a fluence rate of 130 μmol m^−2^ s^−1^, unless otherwise specified. For soil-grown Arabidopsis, Arabidopsis seedlings were geminated on ½ MS plates, and 14-day-old seedlings were transferred to soil (Fafard II) and grown at 22 °C in 16 hour light/8 hour dark cycles with light fluence rate of 135 μmol m^−2^s^−1^ unless otherwise specified.

### Measurement of circadian rhythms and flowering time

The decoy and control *CCA1p::Luciferase* seeds were grown on ½ MS plates with or without 15 μg/ml ammonium glufosinate (Santa Cruz Biotechnology, cat# 77182-82-2). Seven day-old T1 transgenic seedlings were arrayed on ½ MS in a 10×10 grid on a 100 mm square dish, then treated with 5 mM D-luciferin (Cayman Chemical Company, cat# 115144-35-9) dissolved in 0.01% Triton X-100. Seedlings were imaged at 22 °C under constant white light provided by two LED light panels (Heliospectra L1) with light fluence rate of 21 μmol m^−2^ s^−1^. The imaging regime is as follows: each hour lights are turned off for two minutes, then an image is collected using a five minute exposure on an Andor iKon-M CCD camera; lights remain off for one minute after the exposure is completed, and then lights return to the normal lighting regime. The CCD camera was controlled using Micromanager(74), using the following settings: binning of 2, pre-amp gain of 2, and a 0.05 MHz readout mode. Using this set-up, 400 seedlings are simultaneously imaged across four plates. Images are acquired each hour for approximately six and a half days. Data collected between the first dawn of constant light and the dawn of the sixth day were used for analyses.

The mean intensity of each seedling at each time point was calculated using ImageJ(75). The calculated values were imported into the Biological Rhythms Analysis Software System (BRASS) for analysis. The Fast Fourier Transform Non-linear Least Squares (FFT-NLLS) algorithm(76) (http://www.amillar.org/Downloads.html) was used to calculate period and relative amplitude of the rhythms from each individual seedling.

Following circadian analysis, seedlings were transferred to soil and grown in inductive conditions, 22 °C in 16 hour light/8 hour dark cycles with light fluence rate of 135 μmol m^−2^ s^−1^. Plants were monitored daily for flowering status, and dates when each individual reached 1 cm inflorescence height, 10 cm inflorescence height, and showed the first open flower bud were recorded. Additionally, the leaf number for each plant was counted when the inflorescence reached 1 cm.

### Quantification of protein expression in decoy transgenic lines

To determine the relative protein expression levels of decoy transgenic lines, leaf tissue from T1 decoy lines were harvested directly following the completion of flowering time experiments (after the inflorescence stem reached 10 cm). The tissue was frozen and ground in liquid nitrogen. Total protein was extracted from ground powder in urea extraction buffer (8 M Urea, 75 mM NaCl, 50 mM Tris-HCL, pH 8.2), then quantified using a BCA Assay kit (Thermo Scientific, cat#23225). Approximately equal amounts of protein were diluted with a modified Laemmli sample buffer (50 mM Tris-HCl, pH 6.8, 100 mM dithiothreitol (DTT), 8% SDS, 10% glycerol, 0.1% bromophenol blue) for western blot analyses. FLAG-His-tagged proteins were detected with mouse anti-FLAG antibody (Sigma-Aldrich, cat# F3165) and α-tubulin (loading control) was detected with anti-α-tubulin (Sigma-Aldrich, cat# T9026) antibody.

### Immunoprecipitation and mass spectrometry analysis (IP-MS)

For the IP-MS of the ZTL decoy harvested at six hours after dusk (ZT18), the T1 decoy transgenic lines (in a Col-0 background) along with control Col-0 and *35S::FLAG-His-GFP* were grown on ½ MS plates with or without 15 mg/ml ammonium glufosinate (Santa Cruz Biotechnology). Seven day-old T1 transgenic lines were transferred to soil and grown under 16 hour light/8 hour dark conditions at 22 °C for 3 weeks. Prior to harvest, plants were entrained in 12 hour light/12 hour dark with a light fluence rate of 165 μmol m^−2^ s^−1^ at 22 °C for 1 week. Approximately 20 mature leaves from the ZTL decoy lines were harvested at six hours after dusk (ZT18), and an equal number of leaves were harvested from the two control lines at four time points under appropriate light conditions (three hours before dusk in the light, three hours after dusk in the dark, six hours after dusk in the dark, and three hours after subjective dusk in the light; ZT9, 15, and 18 under LD conditions, and ZT15 under LL conditions) and combined prior to further processing. All tissue samples were ground in liquid nitrogen using Mixer Mill MM400 system (Retsch). The immunoprecipitation was done as described previously(72, 77, 78) with modifications for one-step IP. Briefly, protein from 2 ml tissue powder was extracted with sonication in SII buffer (100 mM sodium phosphate pH 8.0, 150 mM NaCl, 5 mM EDTA, 0.1% Triton X-100) with complete™ EDTA-free Protease Inhibitor Cocktail (Roche, cat# 11873580001), 1 mM phenylmethylsplfonyl fluoride (PMSF), and PhosSTOP tablet (Roche, cat# 04906845001). The anti-FLAG antibodies were cross-linked to Dynabeads^®^ M-270 Epoxy (Thermo Fisher Scientific, cat# 14311D) for immunoprecipitation. Immunoprecipitation was performed by incubation with beads at 4 °C for 2h on a tube rocker. The beads were then washed with SII buffer three times, 25 mM ammonium bicarbonate three times, and then 10 mM ammonium bicarbonate twice before being subjected to trypsin digestion (0.5 μg, Promega, cat# V5113) at 37 °C overnight. We vacuum dried the digested peptides using a SpeedVac and then dissolved them in 5% formic acid/0.1% trifluoroacetic acid (TFA). The protein concentration was determined by nanodrop measurement (A260/A280)(Thermo Scientific Nanodrop 2000 UV-Vis Spectrophotometer). An aliquot of each sample was then further diluted with 0.1% TFA to 0.1μg/μl. 0.5μg (5μl) and injected for LC-MS/MS analysis at the Keck MS & Proteomics Resource Laboratory at Yale University.

LC-MS/MS analysis was performed on a Thermo Scientific Orbitrap Elite mass spectrometer equipped with a Waters nanoAcquity UPLC system utilizing a binary solvent system (Buffer A: 0.1% formic acid; Buffer B: 0.1% formic acid in acetonitrile). Trapping was performed at 5μl/min, 97% Buffer A for 3 min using a Waters Symmetry^®^ C18 180μm × 20mm trap column. Peptides were separated using an ACQUITY UPLC PST (BEH) C18 nanoACQUITY Column 1.7 μm, 75 μm × 250 mm (37°C) and eluted at 300 nl/min with the following gradient: 3% buffer B at initial conditions; 5% B at 3 minutes; 35% B at 140 minutes; 50% B at 155 minutes; 85% B at 160-165 min; then returned to initial conditions at 166 minutes. MS were acquired in the Orbitrap in profile mode over the 300-1,700 m/z range using 1 microscan, 30,000 resolution, AGC target of 1E6, and a full max ion time of 50 ms. Up to 15 MS/MS were collected per MS scan using collision induced dissociation (CID) on species with an intensity threshold of 5,000 and charge states 2 and above. Data dependent MS/MS were acquired in centroid mode in the ion trap using 1 microscan, AGC target of 2E4, full max IT of 100 ms, 2.0 m/z isolation window, and normalized collision energy of 35. Dynamic exclusion was enabled with a repeat count of 1, repeat duration of 30s, exclusion list size of 500, and exclusion duration of 60s.

After using the Mascot Distiller program to generate Mascot compatible files, the MS/MS spectra were searched in-house using the Mascot algorithm (version 2.4.0)(79). The data were searched against the Protein SwissProt_2016_05.fasta *Arabidopsis thaliana* database and the significance threshold *p* < 0.5, allowing for oxidation (M), Phosphorylation (STY), and ubiquitination (diGLY-K) as variable modifications. Peptide mass tolerance was set to 10 ppm, the MS/MS fragment tolerance was set to 0.5 Da, and the maximum number of missed cleavages by trypsin was set to two. Normal and decoy database searches were run to determine the false discovery rates. The confidence level was set to 95%. The peptide and protein information is listed in Table S4.

For the IP-MS experiments of the FKF1 and LKP2 decoys, the T2 decoy transgenic lines along with parental *CCA1p::Luciferase* lines and *35S::FLAG-His-GFP* plants were grown on soil in inductive conditions (16 hour light/8 hour dark at 22 °C) for 3 weeks prior to collection at two hours after dusk in long day conditions (ZT18). The immunoprecipitation was done as described previously (72, 77, 78). The immunoprecipitated proteins were digested with trypsin, and the samples were analyzed by LC-MS/MS using a Q-Exactive mass spectrometer (Thermo Scientific) as described previously(79, 80). The mass spectrometry data were processed as described above, except the search parameters were optimized to the mass tolerance of fragment ions with 50 ppm for precursor ions and 0.01 Da for fragments to fit the sensitivity of Q-Exactive mass spectrometer. The peptide and protein information is listed in Table S5.

For the IP-MS of the ZTL decoy time course experiments, we analyzed two biological replicates from the T1 ZTL decoy transgenic line in the *CCA1p::Luciferase* background. The T1 seedlings were selected on ½ MS plates and then grown on soil 16 hour light/8 hour dark at 22 °C. Seven day old seedlings were then transferred to new ½ MS plates without selection for circadian period analysis, as described above.

Then, plants were transferred to soil and grown under 16 hour light/8 hour dark conditions at 22 °C for 2 weeks. The plants were entrained in 12 hour light/12 hour dark with a light fluence rate of 165 μmol m^−2^ s^−1^ at 22 °C for 1 week prior harvest. Approximately 20 leaves from each genotype were harvested at three hours prior to dusk (ZT9, collected in light) and three hours after dusk (ZT15, collected in dark) in 12 hour light/12 hour dark conditions to compare the interacting dynamics of ZTL. At dawn (ZT0), the plants were transferred to continuous light conditions and leaf tissue was harvested at three hours after subjective dusk(ZT15 in LL conditions) to compare to the three hours after dusk (ZT15 in LD) samples.

We pooled parental lines (*CCA1p::Luciferase*) and 35S::FLAG-His-GFP harvested at each time point as controls for IP experiments. A two-step IP protocol was performed as described in the LKP2 and FKF1 IP experiments. The LC-MS/MS analyses were used an Orbitrap Elite mass spectrometer (Thermo Fisher Scientific) and the data were analyzed with MASCOT as described above. The search parameters were same as in the ZTL decoy section described above except for the oxidation (M) peptide modification. The peptide and protein information is listed in Table S6.

### Yeast two-hybrid assays

Yeast two-hybrid assays were performed according to the Yeast Protocol Handbook (Clontech, cat# P3024). Briefly, CHE and TOC1 coding sequences in pENTR/D-TOPO vectors were recombined into the pGBKT7-GW destination vector (Gateway compatible pGBKT7 vector). This resulted in a translational fusion of CHE and TOC1 to the GAL4 DNA-binding domain (GAL4-BD)(77). These constructs were transformed into the yeast Y187 strain. The full-length, LOV domain (a.a. 1-196), Kelch domain (a.a. 246-609), and decoy (the LOV domain fused to the Kelch domain) ZTL coding sequences in pENTR/D-TOPO vectors were recombined into the pGADT7-GW vector (Gateway compatible pGADT7 vector), resulting in a translational fusion to the GAL4 activation domain (GAL4-AD)(77). These were transformed into the yeast AH109 strain. To test protein-protein interactions, diploid yeast was generated by yeast mating of Y187 and AH109 strains bearing pGBKT7 and pGADT7 vectors, respectively, and tested on synthetic drop-out (SD)/-Leu-Trp and SD/-Leu-Trp-His plates. The empty pGBKT7-GW and pGADT7-GW vectors were included as negative controls.

### Co-immunoprecipitation (CO-IP) and *in vivo* ubiquitination assays using HEK293T cells

The full-length (FL) or decoy ZTL (in the pENTR/D-TOPO vectors) was recombined into pEZYflag destination vector to generate translational fusions to the FLAG affinity tag. CHE (in the pENTR/D-TOPO vectors) was recombined into the pEZYegfp destination vector to generate a CHE translational fusion to GFP(81). Approximately 2×10^6^ HEK293T cells in 60 mm petri-dishes were transfected with pEZYflag-ZTL-FL, pEZYflag-ZTL-decoy, pEZYegfp-CHE, and/or pEZYegfp vectors using lipofectamine 2000 (Thermo Fisher Scientific, cat# 11668027) for 24 hours according to the manufacture’s protocol. Prior to harvest, the cells were treated with 30 μM of MG132 (PEPTIDE INSTITUTE INC, cat# 3175-v) for 6 hours. The cells were lysed by sonication in RIPA buffer (Sigma-Aldrich, cat# R0278) with cOmplete™ EDTA-free Protease Inhibitor Cocktail. The co-IP was performed by incubation of cell lysate with anti-FLAG antibody conjugated to SureBeads™ Protein G Magnetic Beads (Bio-Rad, cat# 161-4023) at 4°C for 1 hour. The protein on the beads was washed with SII buffer (100 mM sodium phosphate (pH 8.0), 150 mM NaCl, 5 mM EDTA, 0.1% Triton X-100) three times and eluted by boiling in Laemmli sample buffer for subsequent western blot analyses.

For the *in vivo* ubiquitination assays, approximately 2 × 10^6^ HEK293T cells in 60 mm petri-dishes were transfected with pEZYegfp-CHE, pEZYegfp, pEZYflag-ZTL-FL and/or pEZYflag-ZTL-decoy vectors using lipofectamine 2000 (Thermo Fisher Scientific, cat# 11668027) for 24 hours and then treated with 30 μM of MG132 (PEPTIDE INSTITUTE INC, cat# 3175-v) or DMSO vehicle for 6 hours. The cells were lysed in urea buffer (8 M urea, 50 mM Na-phosphate pH 8.0, 150 mM NaCl, 1mM DTT, cOmplete™ EDTA-free Protease Inhibitor Cocktail) for western blot analyses. For the deubiquitination of GFP-CHE with USP2cc, the cell lysate from the pEZYegfp-CHE and pEZYflag-ZTL-FL transfected and MG-132 treated cell lysate was further used. The cell lysate was diluted 3-fold with 50 mM Na-phosphate and incubated with 2 μg of USP2cc (Sigma-Aldrich, cat# U6653) at 37 °C for 2h. The mouse anti-FLAG and rabbit anti-GFP (Abcam, cat# Ab-290) antibodies were used to detect FLAG- or GFP-fused proteins, respectively.

### Measurement of mRNA and protein levels in CHE-GFP lines

*35S::CHE-GFP* lines were grown on ½ MS plates and entrained in 12 hours of light followed by 12 hours of dark with light fluence rate of 165 μmol m^−2^ s^−1^ at 22 °C for 12 days and then shifted to constant light (LL), constant dark (DD), or kept in 12 hour light/12 hour dark (LD) for 48 hours prior to harvest. For western blotting, the protein was extracted from ground seedlings in 8 M urea buffer (8 M urea, 50 mM Tris pH 8.2, 75 mM NaCl, cOmplete™ EDTA-free Protease Inhibitor Cocktail (Roche, cat# 11836170001)) and quantified with a Pierce™ BCA Protein Assay Kit (Thermos Fisher Scientific, cat# 23225). Approximately 40 μg of total protein was separated on 10% SDS-PAGE for western blot analyses. CHE-GFP protein levels were detected with anti-GFP antibody, and tubulin was detected with anti-α-tubulin antibody (Sigma-Aldrich, cat# T9026). Quantification of signal intensity was performed with Gel Doc™ XR+ System (Bio-Rad) and analyzed with Image Lab software (Bio-Rad). To cross-compare the protein levels among different blots, the ZT9 samples from LL, DD, and LD were included in each blot. The relative protein levels were calculated by average of log_2_ [(CHE-GFP / α-tubulin) _time point A_, _condition X_ / (CHE-GFP / α-tubulin) _ZT9_, _LD_] from three biological replicates.

For quantitative real-time RT-PCR (qRT-PCR), total RNA was extracted from leaf tissue using the RNeasy Plant Mini Kit and treated with RNase-Free DNase (Qiagen, cat# 74904 and 79254) following the manufacturers’ protocols. cDNA was prepared from 100 ng total RNA using iScript™ Reverse Transcription Supermix (Bio-Rad, cat# 1708841) and diluted ten-fold then used directly as the template for the PCR reaction. The qRT-PCR was performed with 4 μl of diluted cDNA and 500 nM primers listed in Table S7 using iTaq™ Universal SYBR^®^ Green Supermix (Bio-Rad, cat# 1725121) with the CFX384 Touch™ Real-Time PCR Detection System (Bio-Rad). The qRT-PCR reactions started with a denaturation step at 95°C for 3 min, followed by 40 cycles of denaturation at 95 °C for 10 sec, primer annealing at 55 °C for 10 sec, and primer extension at 72 °C for 30 sec. *IPP2* (AT3G02780) was used as an internal control. The relative expression of CHE-GFP represents means of log_2_ (2^−ΔΔCT^) from three biological replicates, in which ΔΔCT = (C_T_ of *IPP2* - C_T_ of CHE-GFP) _time point_ − (C_T_ of *IPP2* - C_T_ of CHE-GFP) _ZT9_.

### Cell-free degradation assay

A cell-free degradation assay was done as described previously(24) with modifications as noted below. Frozen 14-day-old seedlings grown under 12 hour light/12 hour dark cycles with a light fluence rate of 165 μmol m^−2^ s^−1^ at 22 °C were collected at four hours after dusk (ZT16) and ground in liquid nitrogen. Powder was re-suspended with ice-cold assay buffer (50 mM Tris-HCl pH 7.4, 100 mM NaCl, 10 mM MgCl_2_, 5 mM DTT, 5 mM ATP, 0.5% Triton-X 100). The plant slurry was split into two tubes and incubated at room temperature with 50 μM MG132 or DMSO vehicle. An equal volume of sample was taken out at designated time points and mixed with Laemmli sample buffers to stop the reactions. All assay steps were carried out in the dark. After boiling for 10 min, the samples were clarified by centrifugation for 10 min at 16000 g. SDS-PAGE was performed on clarified samples for western blotting. CHE-GFP and tubulin were detected with anti-GFP antibody and anti-α-tubulin antibodies, respectively, as described above.

## Acknowledgments

We would like to thank Dr. Dmitri Nusinow and the Keck Proteomics Facility at Yale for materials and assistance with proteomics. We would also like to thank Elan Sherazee, Suyuna Eng Ren, Marshall Deline, Denise George, and Antonette Lestelle for technical and administrative support. We would like to thank Dr. Wei Liu, Dr. Man-Wah Li, Dr. Vivian Irish, and Dr. Adam Saffer for helpful comments on the manuscript.

## Contributions

JMG and EJB conceived of the decoy method. JMG, JPP, CML and SK conceived of the idea and the experiments for CHE. CML, AF, CA, KW, and JMG designed and performed the experiments and experimental analyses. CML, AF, and JMG wrote the manuscript.

## Grant numbers

JMG: NSF EAGER #1548538
CML: Anderson Fellowship, Brown Fellowship
AF: NIH T32 GM007499, Gruber Foundation, NSF GRFP DGE-1122492
SK: NIH R01 GM056006, R01 GM067837, RC2 GM092412
EJB: New Scholar Award from the Ellison Medical Foundation and a Hellman Fellowship
JPP: NIH R01 GM056006

